# Leveraging CRISPR-Cas13d in an inducible knockdown system to interrogate *Drosophila* germ granule mRNAs

**DOI:** 10.1101/2024.09.09.611872

**Authors:** Zoya A. Gauhar, Aaron J. Duthoy, Seema Chatterjee, Rodrigo Berber-Pulido, Cameron Myhrvold, Elizabeth R. Gavis

## Abstract

Ribonucleoprotein (RNP) germ granules are hallmarks of germ cells across the animal kingdom and are thought to be hubs for post-transcriptional regulation that promote formation of the germ cell precursors. While numerous RNAs are associated with germ granules in *Drosophila*, the functions of many in germline development are poorly understood. Current methods for RNA knockdown, such as RNAi, do not allow local depletion of transcripts such as those found in the germ granules. We leveraged CRISPR-Cas13 to create a subcellular RNA knockdown system and tested it on two mRNAs, *nanos (nanos)* and *sarah (sra)*, whose abundance in germ granules differs. Because Cas13 has both *cis* and *trans* cleavage activities, we evaluated the effect of target abundance on off-target RNA depletion. We show on and off-target RNA depletion is coupled when targeting the more abundant *nanos* germ granule transcripts. Off-target RNA knockdown is less potent when the system is used for less abundant *sra* transcripts. When *sra* is knocked down in germ granules, we observed an increase in the calcium indicator GCaMP at the posterior and defective primordial germ cell migration, consistent with *sra* encoding a negative regulator of calcium signaling. In sum, we report an *in vivo* Cas13-based system for subcellular knockdown, evaluate its feasibility, and uncover a novel function for *sra* germ granule transcripts in promoting germline development.

## INTRODUCTION

Early in the development of sexually reproducing organisms, cells make a key decision between germline and somatic fates. While the mechanisms and timing that govern this decision vary across organisms, certain features of germline development are conserved. Among these are germ granules, ribonucleoprotein (RNP) granules that are found in germ cells throughout the animal kingdom [1, 2]. Like many types of RNP granules, germ granules are biomolecular condensates composed of RNAs and RNA-binding proteins held together through networks of RNA-RNA, protein-protein, and RNA-protein interactions [3]. As such, they provide compartments for post-transcriptional regulation.

In some animals, including *Drosophila*, germ granules contain maternal determinants necessary for germline formation [1]. In *Drosophila*, the germ granules reside at the posterior pole of the embryo, as components of a maternally derived cytoplasm called the germ plasm. During embryogenesis, germ granules promote budding of the germline progenitors, called pole cells, from the posterior, and are subsequently incorporated into the pole cells [4]. Within the germ granules, mRNAs are protected from the widespread degradation of other maternal transcripts that occurs during the maternal-zygotic-transition (MZT) [5–7]. Thus, the germ granules provide a supply of maternally-derived mRNAs to the developing pole cells, which are transcriptionally quiescent until gastrulation [8, 9]. These transcripts include *polar granule component* (*pgc*) and *nanos*, which have been shown to play roles in pole cell development including migration to the gonad [10], as well as *Cyclin B* (*Cyc*B). Numerous additional transcripts are likely to associate with the *Drosophila* germ granules based on in situ hybridization studies [11–14], but their roles in promoting germline development remain unknown.

Interrogating the functions of RNAs with complex and dynamic subcellular localizations such as those found in germ granules requires strategies that generate robust subcellular RNA knockdown. Current methods, such as RNA interference (RNAi), are difficult to employ locally in an inducible and temporally regulated manner. Even when expressed under the control of tissue-specific promoters, the RNAi machinery is not readily targeted to specific subcellular regions, limiting its utility for studying transcripts with distinct localizations. Moreover, many mRNAs are enriched rather than exclusively localized to subcellular domains. Indeed, germ granule transcripts are also found at lower concentrations throughout the somatic region of the embryo, where they may play other roles [15–17]. While previous studies have relied on RNAi to investigate the functions of germ granule transcripts, confounding effects on the somatic lineages made it difficult to conclude if the observed defects in germ cell development were due to effects on the germ cells themselves, or were rather influenced by effects on somatic cells that also help promote germline formation [18, 19]. To this end, we leveraged the CRISPR-Cas13 system in an *in vivo* knockdown platform for targeting subcellularly localized RNAs, using *Drosophila* germ granule transcripts as a model (Fig. 1A). Cas13 is an RNA-guided RNA nuclease that can easily be localized to desired subcellular locations, providing a means for local knockdown of target RNAs.

**Figure 1:**
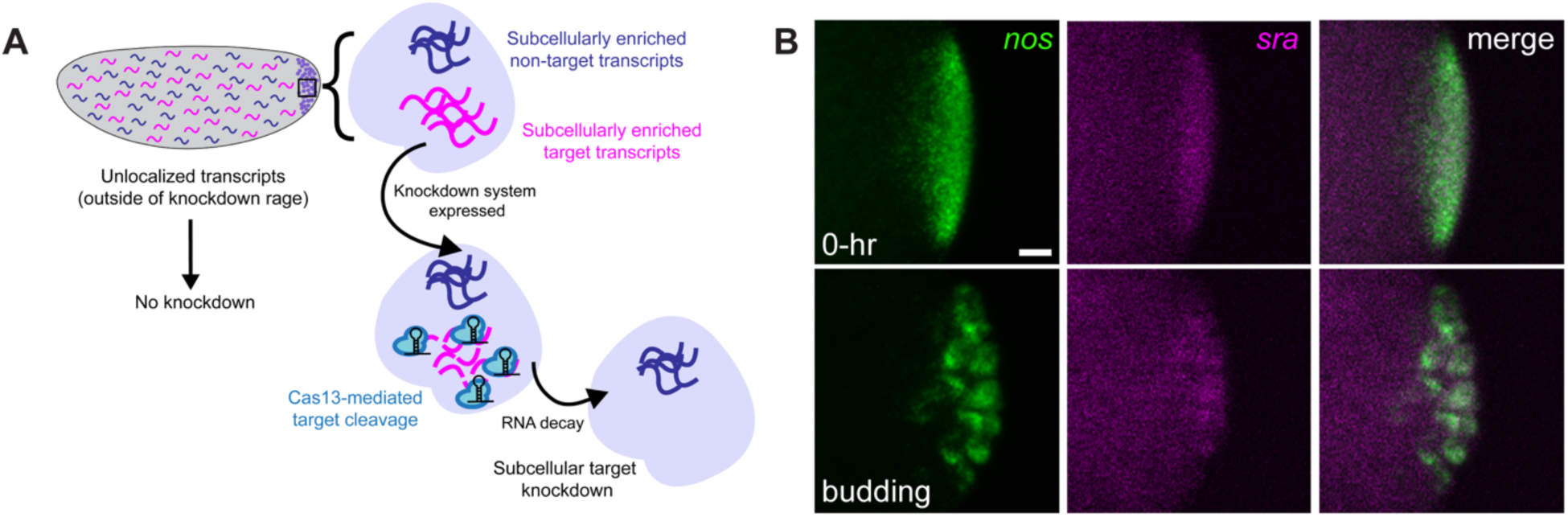
A Cas13-based method for localized RNA knockdown. (A) Strategy for Cas13-mediated knockdown of germ granule transcripts. Spatial control over degradation of a target mRNA (magenta) is conferred by localizing Cas13 protein synthesis to germ granules. (B) Summed slice intensity projections of *nanos* (green) and *sra* (magenta) mRNAs detected by smFISH in 0 hour old and pole-bud stage embryos. *sra* and *nanos* are colocalized in germ granules through pole cell formation, with *sra* less enriched than *nanos*. Scale bar = 10 µm.

Similar to other CRISPR systems, Cas13 effectors can be co-expressed with CRISPR-RNAs (crRNAs) targeting RNAs of interest for cleavage, making them strong candidates for knockdown applications. Cas13 is activated upon crRNA-target binding, and cleaves the target [20]. Following its activation during these *cis* cleavage events, Cas13 remains active and can cleave other target and non-target molecules in *trans* cleavage events [21, 22]. Cas13d, one of the smallest Cas13 effectors, has been previously used in RNA knockdown applications, with conflicting results. Initial studies using cultured mammalian cells suggested Cas13d can confer highly specific target reduction, with limited off-target effects [23]. In subsequent applications, off-target knockdown was frequent and more severe when Cas13d was used to target highly abundant transcripts of interest or in certain cell types [24–27]. In these studies, the off-target activity was attributed to *trans* cleavage events. This variability has also been observed with Cas13d-mediated knockdown *in vivo*. When the Cas13d machinery was injected into zebrafish and other eukaryotic embryos, strong target RNA knockdown with limited off-target effects was observed using RNA-seq and dual reporter assays, respectively [28]. In *Drosophila*, transgenically expressed Cas13d caused toxicity in flies due to off-target RNA degradation in one knockdown application [29], yet caused no toxicity in another [30]. While *trans* cleavage events were not directly attributed to the off-target activity observed, discrepancies between studies may be explained by Cas13’s intrinsic *trans* cleavage activity. These findings highlight the need to monitor potential off-target effects in a Cas13-based knockdown platform in order to determine what specific applications Cas13d may be most amenable to *in vivo*. For subcellular RNAs, unwanted off-target activity could affect RNAs outside of the subcellular location of interest, as well as other non-target RNAs within the targeted region, adding another layer of complexity (Fig. 1A). As germ granules house many different RNAs, they are an ideal test-case to evaluate the feasibility of a local Cas13d-based RNA knockdown platform.

Here, we applied Cas13d in an RNA knockdown system targeting two germ granule targets: *nanos*, which is essential for germline development, and *sarah* (*sra*), which encodes a regulator of calcineurin [31] and whose function in pole cell formation or development is unknown. As these targets are enriched in germ granules to different extents, we evaluated the effect of target abundance on off-target activity, a factor previously shown to affect *trans* cleavage. Using single molecule fluorescent in situ hybridization (smFISH), we show that on and off-target RNA depletion at the posterior of the embryo is coupled when we target highly enriched *nanos* transcripts. Off-target RNA reduction was less apparent for less abundant *sra* transcripts. By contrast, whereas *nanos* was depleted specifically in the germ plasm, *sra* was depleted throughout the embryo. Cas13-mediated knockdown of *sra* did, however, result in enrichment of the calcium indicator GCaMP in pole cells prior to their migration, consistent with Sra protein function. Depletion of *sra* also led to pole cell migration defects, suggesting a novel role for *sra* in promoting germline development. Together, while our results highlight considerations around transcript abundance, we show that Cas13d can be used for subcellular knockdown.

## RESULTS

### Identifying differentially enriched mRNA targets

To select mRNA targets, we consolidated a list of transcripts that appear to colocalize with germ granules based on previous studies and publicly available in situ hybridization data to select targets from [11, 13, 14, 32]. From these data, 25 transcripts have a distribution characteristic of germ granules (Table S1). From this list, we selected *nanos*, whose localization to the germ granules is essential for germline development [33, 34] and *sra*, which encodes a regulator of calcineurin in the calcium signaling pathway [31]. While *sra* has been implicated in *Drosophila* meiosis, ovulation, and plasticity in the neuromuscular junction [31, 35, 36], whether its germ granule localization is functionally significant is unknown, making it an ideal candidate for localized knockdown.

In situ hybridization analysis detected *sra* at the posterior of the early embryo, although it is visibly less enriched than *nanos* as compared to the bulk cytoplasm (Supplementary Fig. 1A). Moreover, *sra* accumulates around budding pole cell nuclei, a distribution characteristic of germ granule mRNAs (Supplementary Fig. 1A). Using single molecule fluorescence *in situ* hybrization (smFISH), we confirmed that *sra* transcripts are indeed weakly enriched in germ granules in the early embryos, using *nanos* mRNA as a germ granule marker (Fig. 1B). Thus, we conclude that *sra* is a germ granule localized transcript that is less abundant in the germ granules compared to *nanos*.

### Generating transgenic *Drosophila* strains for Cas13-mediated knockdown

To increase the likelihood of knockdown, we designed CRISPR arrays containing three crRNAs targeting different regions of either *nanos* or *sra* transcripts (Supplementary Fig. 1B, C). Each array was designed such that Cas13d’s self-processing activity would produce three individual crRNAs from each array. Transgenic flies expressing each array ubiquitously under the U6 promoter were generated, and the expression of each pre-processed array was confirmed using qPCR (Supplementary Fig. 1 D, E).

We also generated flies for expression of HA-tagged CasFx, a *Drosophila* codon optimized variant of Cas13d, whose cleavage activity was previously demonstrated in flies [32], under Gal4-UAS control. To limit knockdown to the target RNAs within germ granules, we fused CasFx coding sequences to the *nanos 3′UTR*. The *nanos 3′UTR* serves two purposes: first, it localizes *Cas13* mRNA to germ granules during their assembly, and second, it limits Cas13 protein synthesis to the germ granules beginning in the late stages of oogenesis [32, 37] (Fig. Supplementary 1F). The resulting *UAS-Cas13-nanos3′UTR* transgene was maternally expressed using *mat-tub:Gal4* and Cas13 expression was confirmed by anti-HA Western blotting of ovary extracts (Supplementary Fig. 1G). Anti-HA immunofluorescence showed that Cas13 protein was highly enriched in the embryonic germ plasm and segregated to the pole cells (Fig. 2B).

**Figure 2:**
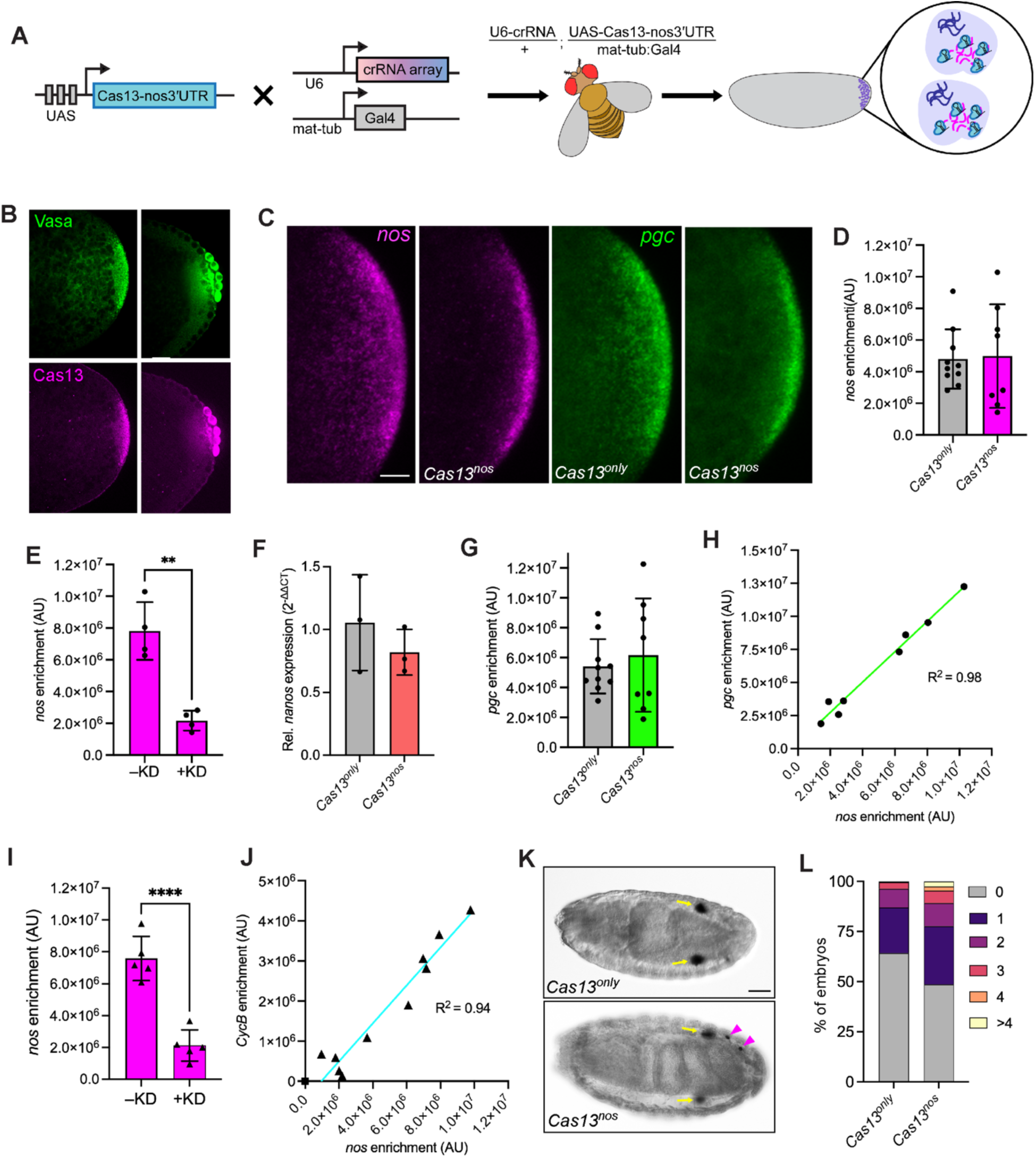
On and off-target activity are coupled in *Cas13^nanos^* embryos. (A) Scheme to produce females co-expressing Cas13 and crRNAs. Cas13-mediated knockdown occurs in eggs produced by these females. (B) Confocal images of Cas13-expressing embryos before (left) and after (right) pole cell formation. The germ plasm is marked with anti-Vasa (green) and Cas13 is detected by anti-HA (magenta) immunofluorescence. (C) Summed slice intensity projections of early *Cas13^only^* and *Cas13^nanos^* embryos, with *nanos* (magenta) and *pgc* (green) detected by smFISH. (D) Quantification of *nanos* enrichment in germ granules for embryos from a single experiment (*n* = 11 embryos for *Cas13^only^*, *n* = 8 embryos for *Cas13^nos^*). (E) Comparison of *nanos* enrichment between *Cas13^nos^* embryos where knockdown occurred and embryos without knockdown based on the bimodally distributed data points. (F) qRT-PCR quantification of *nanos* mRNA in *Cas13^only^* and *Cas13^nos^* embryos. (G) Quantification of *pgc* enrichment in germ granules from the same embryos as in (E). (H) Simple linear regression analysis of the *nanos* and *pgc* levels for individual embryos (*R*^2^ = 0.9931, *y* = 1.562*x* + 686498). (I) Comparison of *nanos* enrichment between *Cas13^nos^* embryos with and without knockdown from a second experimental replicate (*n* = 12 embryos). (J) Simple linear regression analysis of the *nanos* and *CycB* levels for individual embryos (*R*^2^ = 0.9781, *y* = 0.7040*x* + 418194). (K) DIC images of Bownes stage fourteen *Cas13^only^* and *Cas13^nos^* embryos. Anti-Vasa immunohistochemistry was used to detect pole cells. Pole cells populating the gonad are indicated with yellow arrows, “lost” pole cells with magenta arrowheads. (L) Quantification of embryos with the indicated number of lost post cells (*n* = 459 embryos for *Cas13^only^*, *n* = 658 embryos for *Cas13^nos^*). Data points represent individual embryos except in (F) where they represent biological replicates; *\*\*\*\**p<0.0001 and *\*\*\**p<0.001 as determined by an unpaired t-test. Unpaired t-test of ΔΔCt values used to calculate fold-change in (F) showed no significant difference. Scale bars = 10 µm (B) and 50 µm (K).

### On-target and off-target knockdown are coupled for abundant *nanos* target transcripts

To investigate knockdown efficiency, we generated flies expressing Cas13 alone or with the *nanos* crRNA array, and analyzed the embryos they produced (Fig. 2A). We performed smFISH to detect *nanos* in both early embryos expressing the CRISPR-Cas13 machinery targeting *nanos* germ granule transcripts (*Cas13^nos^*; see Methods for nomenclature) and control embryos expressing Cas13 without crRNAs (*Cas13^only^*) (Fig. 2C). Consistently, we observed two populations of embryos, one with reduced enrichment of *nanos* in the germ plasm and the other with enrichment comparable to *Cas13^only^* control embryos (Fig. 2D; replicates in Supplementary Fig. 2A, H). This bimodal distribution may be explained by mosaic expression of the Gal4 driver used to activate transcription of Cas13 in the ovarian nurse cells, causing variation in the amount of Cas13 protein per embryo [38] (Supplementary Fig. 2E). Therefore, to evaluate the extent of knockdown (KD), we conservatively defined KD as being outside the lower bound of the confidence interval for the *Cas13^only^*control. With this criterion, the difference between +KD and -KD is significant (Fig. 2E, I and Supplementary Fig. 2B). qRT-PCR analysis showed that the total amount of *nanos* mRNA was similar in *Cas13^nos^* and *Cas13^only^* embryos and is consistent with selective targeting of germ granule localized *nanos*, which constitutes only 4% of the total [15] (Fig. 2F).

Due to Cas13’s propensity for off-target cleavage events from *trans* cleavage activity, we investigated whether off-target RNA reduction (defined as depletion of RNA not targeted by the crRNA), could be detected in *Cas13^nos^* embryos. We performed smFISH using probes for another germ granule transcript, either *pgc* or *CycB,* as a Non-Target Control (NTCs) (Fig. 2C and Supplementary Fig. 2F). Each germ granule NTC exhibited a bimodal distribution that was similar to that of the target *nanos* mRNA (Fig 2G and Supplementary Fig. 2C, I). Consistently, *nanos* mRNA enrichment showed a linear relationship with NTC levels (Fig. 2H, J and S2D). As a decrease in NTC levels was only observed when *nanos* enrichment also decreased, these data suggest that off-target RNA reduction is coupled with target reduction. This is consistent with what is expected from Cas13d’s *cis* and *trans* cleavage activities in mammalian cells [39].

### Late-stage *Cas13^nos^* embryos have germ cell migration defects

Originating at the posterior of the embryo, the pole cells migrate during gastrulation to populate the gonad [40]. Pole cells lacking *nanos* or *pgc* transcripts frequently fail to reach the gonad [41, 42]. Thus, we next asked if Cas13-mediated knockdown of *nanos* mRNA and secondary depletion of *pgc* mRNA had effects on pole cell migration. Pole cells were visualized in late-stage embryos during the period of gonad formation using immunohistochemical (IHC) staining for the pole marker, Vasa (Vas) (Fig. 2K). Compared to wild-type embryos, *Cas13^nos^* embryos exhibited a greater number of pole cells that failed to reach the gonad (Fig. 2L). Notably, the distribution of *Cas13^nos^*embryos with at least one “lost” pole cell aligns with the bimodal distribution of *nanos* and *pgc* mRNA levels in early embryos. In addition to promoting formation of the germline, posterior enrichment of *nanos* mRNA is essential for embryonic patterning. Embryos with reduced *nanos* at the posterior have fewer than the wild-type 8 abdominal segments [43]. While *Cas13^nos^* larvae did not exhibit a dramatic loss of abdominal segments, a small number of larvae exhibited segmental fusions (Supplementary Fig. 2J). Anterior morphology was similar to wild-type in *Cas13^nos^* larvae. Together, these data suggest that the migration defects observed in *Cas13^nos^* embryos are likely due to combined effects of *nanos* and *pgc* mRNA depletion that is limited to the posterior.

### Targeting of a less highly enriched mRNA

After we did not observe complete knockdown of *nanos,* an abundant germ granule transcript, we next asked if a more complete knockdown of the less enriched *sra* germ granule transcripts could be achieved. We generated transgenic lines expressing an array containing three *sra* crRNAs, and fortuitously isolated one line containing two insertions of the transgene (*sra\*\**). This doubly inserted line expressed more pre-processed *sra* crRNAs than the single insertion (Supplementary Fig. 1E), which allowed us to test if expressing more crRNAs had an effect on target knockdown. Analysis of *sra* by smFISH across three replicate experiments showed that *sra* enrichment was reduced in *Cas13^sra^* and *Cas13^sra**^* embryos as compared to *Cas13^only^* embryos (Fig. 3A, B). Moreover, many of these embryos had enrichment values around 0, which suggests that almost complete knockdown of a low abundance target is more achievable even with a single crRNA insertion. RT-qPCR analysis revealed that total *sra* levels were decreased in *Cas13^sra^* as compared to *Cas13^only^* embryos however (Fig. 3C), indicative of knockdown of *sra* RNA throughout the embryo, not just in the germ granules. In this case, the spatial specificity of knockdown was lost (see Discussion).

**Figure 3:**
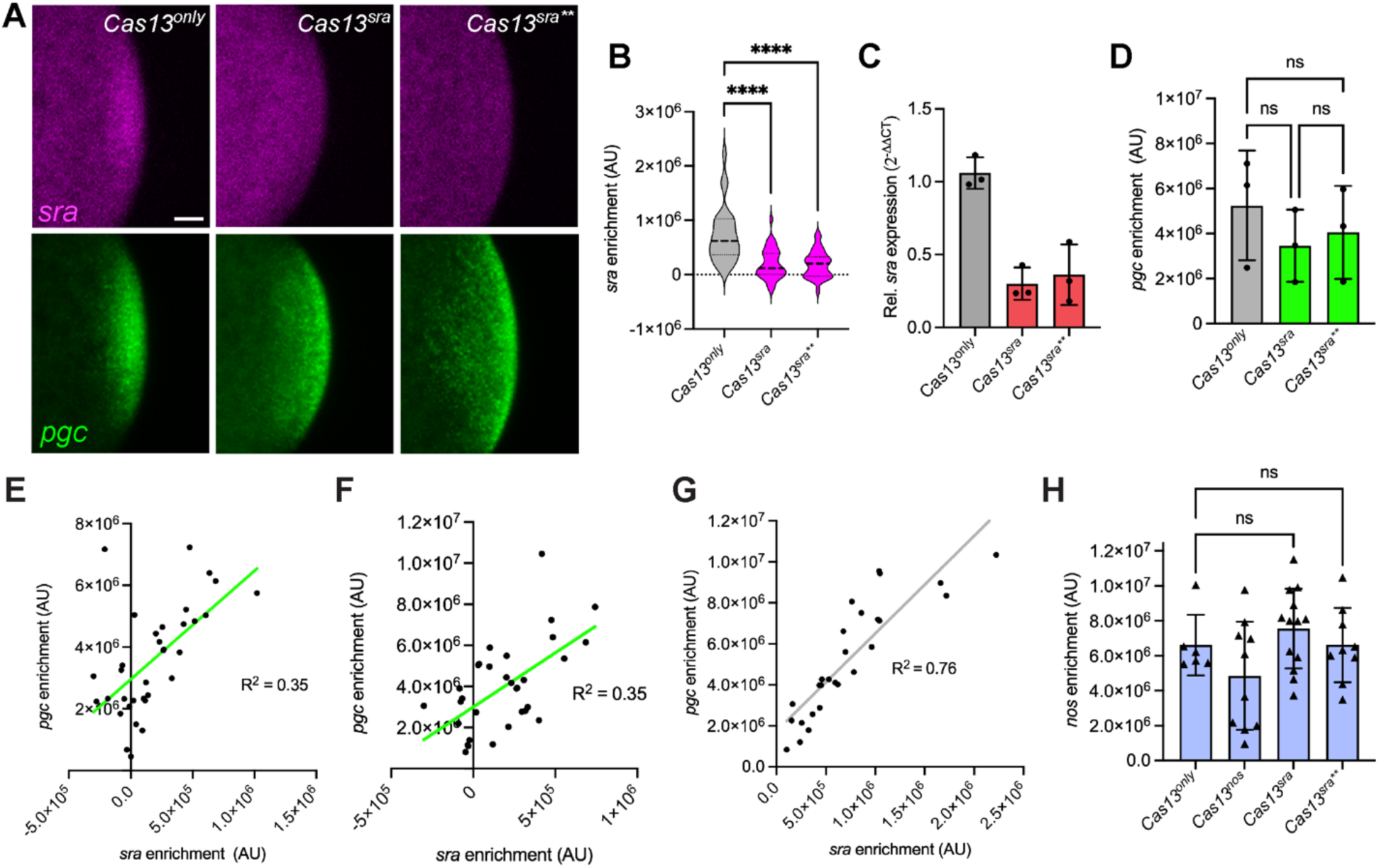
Cas13-mediated knockdown of a lower abundance target is more effective, with less off-target activity. (A) Summed slice intensity projections of early *Cas13^only^*(A), *Cas13^sra^* and *Cas13^sra**^* embryos with *sra* (magenta) and *pgc* (green) detected by smFISH. (B) Violin plot of *sra* enrichment in germ granules for three experimental replicates (*n* = 27 embryos for *Cas13^only^*, *n* = 35 embryos for *Cas13^sra^*, and *n* = 32 embryos for *Cas13^sra**^*). (C) qRT-PCR quantification of *sra* mRNA in *Cas13^only^*, *Cas13^nos^*, and *Cas13^sra**^*embryos. (D) Quantification of mean *pgc* enrichment for the three experimental replicates. (E-G) Simple linear regression of the *sra* and *pgc* enrichment for *Cas13^sra^* (F; *R^2^*=0.3540, *y* = 3.516*x* + 2955538), *Cas13^sra**^* (G; *R^2^* = 0.3511, *y* = 5.278*x* + 2994481), and *Cas13^only^* (H; *R^2^=* 0.7626, *y* = 4.759*x* + 1743067) embryos. Linear regression of *Cas13^only^* shows scatter of *pgc* enrichment intensity in control scenario. (H) Quantification of *nanos* enrichment for the indicated genotypes. Data from the experiment in Fig. 2 I, J and Fig. S2 H, I for *Cas13^only^* and *Cas13^nanos^* embryos are included alongside data from *Cas13^sra^* and *Cas13^sra**^*embryos, which were all collected at the same time to perform *nanos* smFISH. Data points represent individual embryos (B, E-H), biological replicates (C) and means (D); p<0.05 for comparison of ΔΔCt values for *sra* and *sra\*\** to WT that were used to calculate fold changes in (C), **p<0.01, *\*\*\*\**p<0.0001, ns = not significant as determined by ordinary one-way ANOVA and Dunnett’s multiple comparisons test. Scale bar = 10 µm.

### Reduced NTC reduction for a less abundant target

In previous studies, non-target RNA reduction from *trans* cleavage events in cultured cells was less severe when Cas13d was used to target low abundance transcripts [24–26, 39]. To determine if this was also true in our system, we calculated *pgc* enrichment, and found there was no significant difference in *pgc* enrichment between *Cas13^only^*, *Cas13^sra^*, and *Cas13^sra**^* (Fig. 3D). Furthermore, *sra* and *pgc* enrichment were poorly correlated in *Cas13^sra^* and *Cas13^sra**^* embryos, though reduced *pgc* enrichment was only observed when reduced *sra* enrichment was also observed (Fig. 3E, F). Slight variations in *pgc* enrichment occurred even in wild-type embryos (Fig. 3G). Together, these data indicate that indeed, the effects of NTC knockdown in the germ plasm are less pronounced for Cas13-mediated knockdown against a low abundance target.

We wondered if the observed reduction in off-target effects in *Cas13^sra^* and *Cas13^sra**^* affects other germ plasm mRNAs. We performed smFISH in *Cas13^nos^*, *Cas13^sra^*, and *Cas13^sra**^* embryos, and probed for *nanos*. Unlike the bimodal distribution of *nanos* levels observed in *Cas13^nos^*, *nanos* germ plasm levels were not significantly different in *Cas13^sra^*and *Cas13^sra**^* embryos (Fig. 3H). Moreover, even the lowest *nanos* level in *Cas13^sra^* and *Cas13^sra**^* embryos was higher than those of *Cas13^nos^*embryos exhibiting knockdown, suggesting that off-target cleavage events alone are not sufficient to produce knockdown comparable to that achieved with a crRNA for a specific target. Together, these data indicate that our Cas13d-based system confers more specific knockdown against a lower abundance target.

### *sra* promotes pole cell migration

Although *sra* knockdown was not limited to germ granules, *Cas13^sra^*or *Cas13^sra**^* embryos developed without obvious morphological defects, suggesting that depletion had little effect outside the germ plasm. The ability to deplete *sra* mRNAs effectively in germ granules without obvious off-target effects motivated us to investigate a role for *sra* in pole cells. *sra* encodes calcipressin, a negative regulator of calcineurin [31, 44–47]. As a previous study found that the *sra 3′UTR* confers a translational onset prior to pole cell migration [32], we investigated whether Cas13-mediated knockdown of *sra* affects calcium waves occurring in the embryo prior to pole cell migration, using the calcium sensor GCaMP [48] expressed transgenically in the germline (Fig. 4A). Genetic limitations prevented us from introducing a pole cell marker into flies expressing the Cas13 machinery together with GCaMP. Instead, we visualized at GCaMP enrichment at the posterior in embryos where pole cells were morphologically visible. *sra^Cas13^* embryos exhibited a significant increase in GCaMP enrichment specifically in the pole cells prior to migration (Fig. 4B). We then asked if Cas13-mediated knockdown of *sra* transcripts might lead to abnormal pole cell migration. Anti-Vas IHC showed an increased frequency of pole cell migration defects in both *Cas13^sra^* and *Cas13^sra**^* embryos (Fig. 4C, D and Supplementary Fig. 3). Over 10% of *Cas13^sra^* and *Cas13^sra**^* embryos had more than four “lost” pole cells, with some located far from the gonads (Fig. 4B). We cannot rule out the possibility that the migration defect is due in part to knockdown of *sra* in the soma. However, taken together, these data suggest that *sra* germ granule mRNAs may promote pole cell migration during embryogenesis through the calcium signaling pathway.

**Figure 4:**
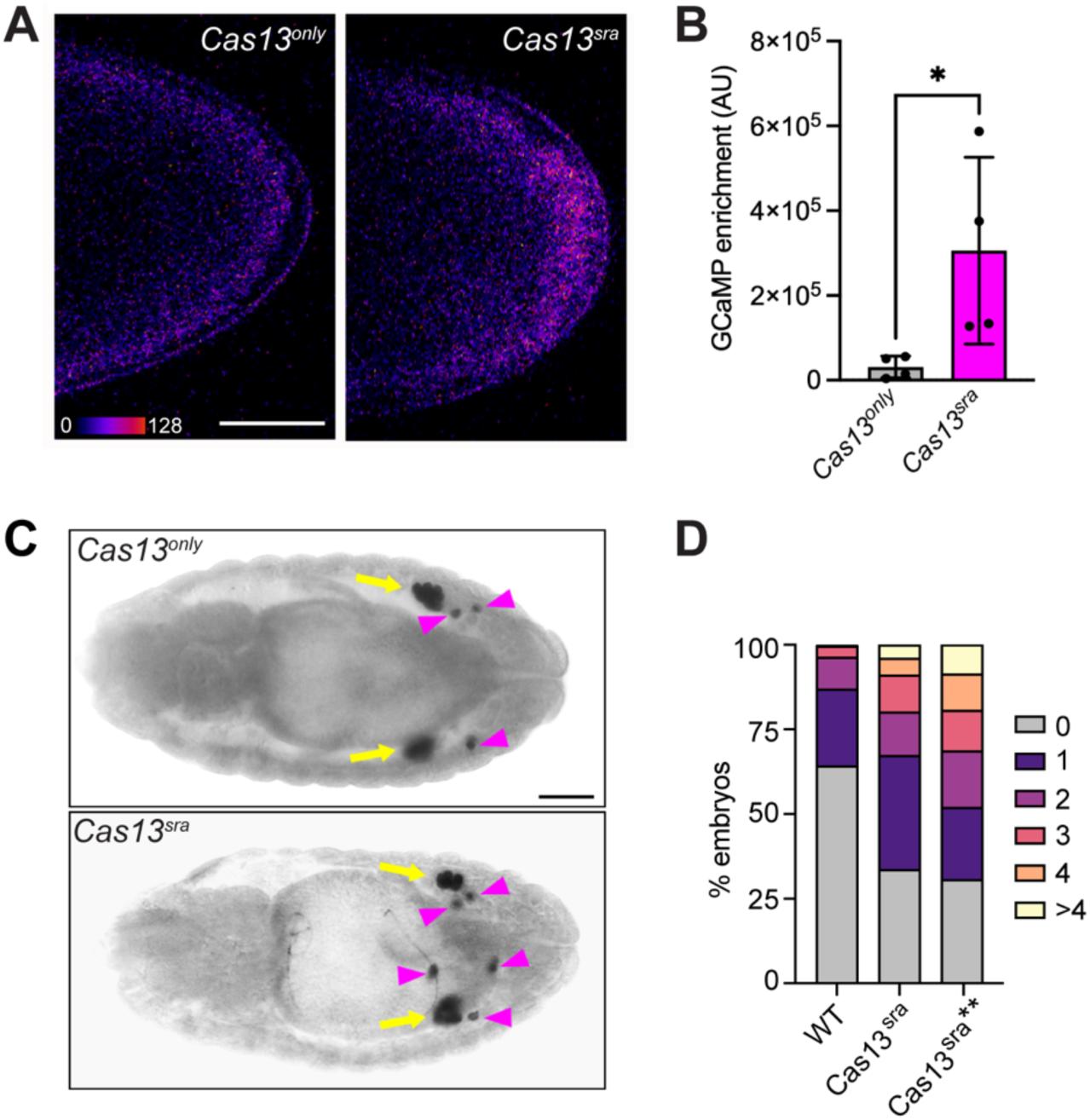
*sra* regulates calcium signaling in pole cells and promotes pole cell migration. (A) Still images from time-lapse imaging of the GCaMP sensor in *Cas13^only^* and *Cas13^sra^* embryos. Single sections through the middle of the embryo are shown. (B) Quantification of GCaMP fluorescence intensity (*n* = 4 embryos for each genotype). (C) DIC images of *Cas13^sra^* and *Cas13^sra**^* Bownes stage 14 embryos. Pole cells were detected with anti-Vasa immunohistochemistry. Black arrows indicate pole cells in gonads, and magenta arrowheads indicate “lost” pole cells. (D) Quantification of embryos with the indicated number of lost pole cells (*n =* 101 embryos for *Cas13^sra^* and *n =* 150 embryos for *Cas13^sra**^*). Data points represent individual embryos; *\**p<0.05 by an unpaired t-test. Scale bars = 50 µm (A) and 100 µm (C).

## DISCUSSION

While germ granules are a common feature of germline development across species, the roles of many germ granule mRNAs have yet to be interrogated due to a lack of methods for probing the functions of subcellularly enriched transcripts. Here, we implement a Cas13d-based RNA knockdown platform to investigate those functions in *Drosophila*. Using this system, we begin to study the feasibility of this strategy and uncover a novel function for one germ granule transcript, *sra*.

### Cas13-based subcellular RNA knockdown

To study complex questions about RNA regulation *in vivo*, the ability to manipulate the expression of individual components is essential for a subcellular mRNA knockdown system. As Cas13 proteins are exogenous to eukaryotic systems, achieving local knockdown using localization tags at the transcript and/or protein level is a promising approach. In addition to localization, temporal control may also be imposed to study particular time periods. Ubiquitously expressing crRNAs, as in our strategy, adds additional flexibility. Previously generated crRNA-expressing animals may be crossed to various localized-Cas13-expressing animals, to deplete the same transcripts in different subcellular domains.

We attempted subcellular RNA knockdown of germ granule enriched mRNAs by targeting Cas13 protein production to *Drosophila* germ granules using the *nanos* 3′UTR, while ubiquitously expressing crRNAs. The *nanos* 3′UTR was also intended to provide temporal control, by initiating localized Cas13 protein synthesis during the late stages of oogenesis [49]. This strategy succeeded for spatially restricted knockdown of *nanos* mRNA, resulting in depletion of *nanos* in germ granules without affecting *nanos* mRNA throughout the somatic region of embryo. It failed, however, to limit *sra* depletion to germ granules, suggesting that there is some amount of Cas13 protein throughout the embryo. Although Cas13 is translated at the posterior, it may diffuse anteriorly prior to pole cell formation similarly to Nanos protein, which forms an anterior-to-posterior gradient. Whereas the amount of Cas13 that permeates the somatic region may be insufficient to target *nanos* mRNA, it may be sufficient to target *sra*, whose concentration in the soma is 5-fold less [17]. In the future, localizing Cas13 by fusing it to a germ granule resident protein such as Oskar should overcome this limitation.

Along with flexibility, limited off-target activity is an essential component of any knockdown system. Given Cas13’s propensity for *trans* cleavage activity, we measured target and NTC mRNA depletion on a per embryo basis using smFISH analysis. Indeed, we observed off-target activity when targeting *nanos* mRNA. While we cannot directly attribute this off-target activity to *trans* cleavage events occurring *in vivo,* the correlation between target and NTC knockdown has also been observed for Cas13d in mammalian cell culture [39]. As target abundance has previously been shown to affect Cas13’s propensity for *trans* cleavage [24–27], we took advantage of *sra*’s weak enrichment at the posterior to evaluate NTC depletion when targeting a less abundant RNA. Indeed, targeting *sra* for knockdown caused more robust on-target depletion together with less off-target effects. Cas13d’s varying propensity for *trans* cleavage activity depending on target abundance may provide one explanation for confounding results in previous *in vivo* applications, which target different RNAs in different cell contexts [28, 30, 50]. Based on our data, we conclude that Cas13d may be better suited for studying low abundance targets *in vivo*. Our results also suggest that *cis* and *trans* cleavage activities can still be coupled when Cas13 is transgenically expressed *in vivo*, which could potentially be useful for applications which require robust local depletion of many different RNA targets at once.

In future applications, *trans* cleavage may be limited using different Cas13 effectors shown to have less *trans* cleavage activity in cell contexts such as Cas13b [24] or high-fidelity Cas13 variants [27], which are engineered to still have robust *cis* cleavage with the absence of *trans* cleavage. Despite limitations due to off-target activity, Cas13d-based RNA knockdown resulted in robust reduction of a less enriched germ granule mRNA, *sra*, which allowed us to begin interrogating its function in the context of germ cell development.

### Interrogating germ granule mRNAs and *sra* function

Although numerous mRNAs are thought to occupy *Drosophila* germ granules [10–13,49], only a small subset have been functionally characterized. As germ granules supply maternally deposited transcripts to the mostly transcriptionally quiescent pole cells [8, 9], it seems likely that other germ granule mRNAs outside of the subset also play roles in promoting germline development. Our finding that Cas13-mediated knockdown of *sra* results in defective pole cell migration supports this idea. As *sra* negatively regulates calcineurin activity [44–47], one potential mechanism for germ granule localized *sra* mRNA to promote germ cell migration is by modulation of calcium signaling. While we cannot rule out a role for *sra* in the soma and we have not been able to directly test the effect of *sra* knockdown in migrating pole cells, our finding that Cas13-mediated *sra* knockdown leads to elevated GCaMP in the pre-migratory pole cells supports this hypothesis. Modulation of calcium signaling has been previously observed to promote cell migration processes [51, 52] and effects on calcium signaling are also well documented in tumor cell migration [53–55]. Polarization of calcium waves promotes convergent extension in amphibians during gastrulation [56–58]. Thus, modulation of calcium fluxes in pole cells by Sra protein may be one developmental consequence of germ granule mRNA localization.

## ACKNOWLEDGEMENTS

We are grateful to M. Wolfner for the *UASp-GCaMP* flies and T. Schüpbach for the Anti-Vas antibody. We thank G. Laevsky and S. Wang in the Princeton Confocal imaging facility, a Nikon center for excellence in the Department of Molecular Biology, for assistance with microscopy, K.Y. Lee for generating the *nanos* riboprobe used in Fig. S1, and A. Hakes and M. Kilwein for technical advice. Research reported in this publication was supported by the National Institute of General Medicine of the National Institutes of Health under grant numbers R35 GM126967 to E.R.G., T32GM007388 to R.B.-P., and by the National Institute of Allergy and Infectious Diseases of the National Institutes of Health under grant number R01 AI182281 to C.M. Funds were also provided by a Princeton University Dean for Research award to E.R.G. and C.M.

## Data Availability Statement

The authors affirm that all data necessary for confirming the conclusions of the article are present within the article, figures, and tables. Strains and plasmids are available upon request.

## Disclaimer

The content is solely the responsibility of the authors and does not necessarily represent the official views of the National Institutes of Health.

## METHODS

### *Drosophila* strains and genetics

The following fly stocks were obtained: *UASp-GCaMP* (gift from M. Wolfner [48]), *maternal-tubulin:Gal4^v37^*(BDSC #7063). For Cas13-mediated knockdown, flies homozygous for both the *maternal-tubulin:Gal4^v37^*driver on chromosome 2 and a crRNA array on chromosome 3 (see below) were crossed to flies homozygous for the *UAS-Cas13-nanos3′UTR* transgene (see below). Progeny are labeled as *Cas13^nos^* or *Cas12^sra^* to indicate to which RNA is targeted. Note that the although abbreviation of *nanos* to *nos* has been discontinued, we have continued to use this abbreviation for brevity. *Cas13^only^* control flies were generated by crossing flies homozygous for the *mat-tub:Gal4* driver to flies homozygous for the *UAS-Cas13-nanos3′UTR* transgene.

### crRNA design

crRNAs were designed to target the most representative isoforms of each target in the 0-2 hour embryo, based on FLYBase high throughput expression data. *nanos-RA* (FLYBase ID:FBtr0083732) and *sra-RA* (FLYBase ID: FBtr0083255) were chosen as targets.

Taking advantage of Cas13’s crRNA processing activity, three unique Cas13d crRNAs were designed per target and expressed as an array. Each guide region was designed using the publicly available Cas13design tool [59, 60]. To generate guide RNAs, *Drosophila* was selected as the model organism, and the associated FlyBASE ID for each target isoform was inputted. Three sequences complementary to unique regions within each target transcript were selected. All guide sequences are shown in Table S2. All predicted guides had scores of above 85%, and regions targeted coding sequences (CDS) were selected over untranslated regions (UTRs) when possible. For compatibility with Cas13d, the direct repeat sequence (Table S2) was added at the 5’ end, followed by a guide sequence to create a complete crRNA. For each target, the three crRNAs were synthesized as an array followed by a terminator sequence (Table S2) with flanking NotI and PstI digest sites (Genewiz).

### *Drosophila* transgene generation

crRNA arrays were cloned into a vector previously used to express Cas13d crRNAs in *Drosophila* [50]. The *nanos* crRNA array was digested using NotI and the blunt cutter MslI. The vector was digested using NotI and PstI digest sites and end-filled using T4 polymerase, prior to ligating the *nanos* crRNA insert into the vector. The *sra* crRNA array was digested using NotI and PstI digest sites, and ligated into NotI and PstI digested vector. crRNA arrays were was inserted at the attP40 site on the second chromosome using the ΦC31 recombinase system.

To generate the *UAS-Cas13-nanos3′UTR* transgene, the *Drosophila* codon optimized Cas13d sequence (CasFX4; [30]) was synthesized, including 3xHA tag just before the first CasFX4 codon (Genewiz), flanked by KpnI and HindIII sites. This sequence was inserted together with a HindIII-NotI fragment containing the *nanos* 3′UTR and 75 bp of 3′ genomic DNA between KpnI and NotI sites of pattB-UASp from which the K10 3′UTR sequences were removed. The *UAS-Cas13-nanos3′UTR* transgene was inserted at the VK0033 site on the third chromosome using the ΦC31 recombinase system.

### In situ hybridization

For histochemical *in situ* hybridization, embryos were collected for 2.5 hrs at 25 ℃. Embryos were dechorinated, fixed, and devitellinized using methanol as previously described [60], and stored in methanol at -20 ℃ for up to two weeks. In situ hybridization was performed as previously described with using digoxigenin-labeled riboprobes [37]. For the *sra* riboprobe, cDNA (#LD15403) from the Drosophila Genomics Resource Center was used as a template.

For smFISH, embryos were collected on yeasted apple juice plates for 2.5 hours at 25 ℃ and staged by nuclear density and morphological features for nuclear cycle 0. Embryos were dechorinated, fixed, hand peeled as previously described [61], dehydrated stepwise into methanol and stored at -20 ℃ for up to one week. smFISH was performed as previously described [60] after rehydrating embryos stepwise into PBST (PBS/0.1% Tween20), and the samples were then mounted in Vectashield. Prior to imaging, embryos were stored overnight at 4 ℃ with the coverslip facedown. For *sra* RNAs, a custom *sra*-570 probe set was generated (Stellaris).

### Immunofluorescence

Embryos were collected for 3 hours at RT and fixed as above. Embryos were rehydrated stepwise into PBST (1× PBS, 0.1% Tween-20), then blocked in BBT (PBST, 0.1% globulin-free BSA) for 2 hours at RT. Primary antibodies diluted 1:500 in BBT were applied overnight at 4 ℃. Embryos were washed in BBT, then incubated for 30 minutes in BBT+2% normal goat serum. Secondary antibodies diluted 1:1000 in BBT+2% were applied serum for 2 hours at RT. After washing in PBST, embryos were mounted in Vectashield. Antibodies: mouse anti-HA (BioLegend clone 16B2); rabbit anti-phospho-Vasa (gift of T. Schüpbach); goat anti-mouse-Alexa 568 and goat anti-rabbit-Alexa 647 (Thermo Fisher Scientific).

### Imaging and image quantification

Confocal imaging for smFISH analysis was performed using a Nikon A1 scanning confocal microscope using GaAsp detectors and a 60 × 1.4 NA oil immersion objective, with the exception of Figure S2A, where a 40 x 1.3 NA oil immersion objective was used. Z-series were captured using a 1.5 µm step-size. Within each experiment, imaging parameters among samples were kept identical. smFISH was quantified by generating “summed slice intensity” projections in FIJI. Enrichment of mRNA in the germ plasm was calculated by measuring the integrated density of the germ plasm within an ROI selected using the germ plasm marker (either *pgc* or *CycB*). The ROI was moved to a region of the cortical cytoplasm to measure a background value, which was then subtracted from the germ plasm value.

Images of Cas13 and Vasa immunofluorescence were acquired on a Nikon W1-SoRa spinning disc confocal microscope using an apo TIRF 60x oil DIC N2 objective.

### Immunohistochemistry

Embryos were collected on yeasted apple juice plates for six hours at 25 ℃, then aged for 20 hours at 18℃. Anti-Vas IHC was performed as previously described [62] to detect pole cells in Bownes staged 14-15 embryos with the following antibodies: 1:500 rabbit anti-phospho-Vasa and 1:500 biotin goat anti-rabbit (Jackson Immuno Research Laboratories). Embryos were mounted in 80% glycerol and pole cells were visualized and counted on a Zeiss compound microscope. Images of representative embryos were captured using a Nikon DsQi-2 microscope, with a 20 X 0.75 NA air objective and DIC optics.

### Larval cuticle preparation

Embryos were collected on yeasted apple juice plates at room temperature over one day, aged for one day, and stored at 4 ℃ for up to one week. Cuticle preparations were performed as previously described [63], mounted in Hoyer’s media, and segments were counted on a Zeiss compound microscope.

### Live-imaging and quantification of GCaMP

For live-imaging, approximately 2.5 hour old embryos were collected on yeasted apple juice plates at 25 ℃, dechorinated, placed onto a #1.5 coverslip using a paintbrush, and covered in a drop of Halocarbon 200 Oil. Live-imaging was performed on a Nikon A1 scanning confocal microscope using GaAsp detectors and a 40 x 1.3 NA oil objective, which was focused to the middle of the embryo, and images were captured every forty seconds. For GCaMP fluorescence quantification, a single frame from each video was selected where the pole cells were in a similar plane. Integrated density was measured for the germ plasm, as well as for a region of the cortical cytoplasm of the same area. Enrichment was calculated by subtracting the germ plasm integrated density from the cortical cytoplasm integrated density.

### RT-qPCR

#### crRNA quantification

Ovaries were dissected from wild-type females or females homozygous for either crRNA transgene, flash frozen in liquid N_2_, and stored at -80 ℃. RNA was extracted from 20 ovaries per sample using the *Quick*-RNA^TM^ Miniprep Plus Kit (Zymo Research). Genomic DNA removal and cDNA generation were performed using the Quantitect Reverse Transcription Kit. In place of RT primer mix, transgene specific primers (Table S3) designed for each crRNA were used to generate cDNAs. qPCR was performed using SYBR Green PCR Master Mix (Thermo Fisher Scientific) and the StepOnePlus Real-Time PCR system (Applied Biosystems) according to manufacturer’s instructions using the same gene specific primers. Three technical replicates were performed per sample. For analysis, technical replicates were averaged, and the mean CT was used to calculate the relative gene expression per sample normalized to the wild-type control by the ΔΔCT method [64], with *rp49* used as the reference gene.

#### nanos and sra quantification

Embryos were collected for 1.5 hours, dechorionated, flash frozen in liquid N_2_, and stored at -80 ℃. RNA was extracted from embryos using RNeasyTM extraction kit (Qiagen). Genomic DNA removal and cDNA generation were performed using the Quantitect Reverse Transcription Kit, using the oligo dT primers provided with the kit. qPCR and data analysis was performed as described above, with three biological and three technical replicates for each sample.

### Immunoblotting

Flash frozen ovaries from either of two independent transgenic *Cas13-nanos3′UTR* lines or with the *mat-tub:GAL4* driver alone were thawed and dissolved in 5 µl of sample buffer (0.125 M Tris-HCl (pH 6.8), 4% SDS, 20% glycerol, 5 M urea, 0.01% bromophenol blue, and 0.1 M DTT) per ovary and boiled for 4 minutes. The samples were run on a 9% SDS-PAGE gel, and immunoblotting was performed as previously described [63] using the following antibodies: 1:1000 rabbit anti-HA (Cell Signaling Technologies #C29F4) and 1:2000 HRP donkey anti-rabbit (GE Healthcare). The blot was imaged using an iBright FL1000 imaging system. Due to an interaction between the HA and tubulin antibodies, the same blot was stripped using Restore Western Blot Stripping Buffer (Thermo Scientific), reprobed with the following antibodies: 1:5000 mouse ascites (Sigma) and 1:1000 HRP sheep anti-mouse (NA931), and reimaged.

