## Supplementary Information for "Leveraging CRISPR-Cas13d in an inducible knockdown system to interrogate *Drosophila* germ granule mRNAs"

Running title: CRISPR-Cas13 knockdown of *Drosophila* germ granule RNAs

### SUPPLEMENTARY TABLES

| |
|---|
| *nanos* |
| *germ cell-less* |
| *polar granule component* |
| *Cyclin B* |
| *Septin 4* |
| *spire* |
| *pumilio* |
| *oo18 RNA-binding protein* |
| *CG11597* |
| *Blastoderm-specific gene 25D* |
| *Tao* |
| *Linking immunity and metabolism* |
| *CG2865* |
| *bruno 1* |
| *CG31998* |
| *Rap GTPase activating protein 1* |
| *Pinocchio* |
| *Sarah* |
| *Discs large 5* |
| *Ras GTPase activating protein 1* |
| *Upstream of N-ras* |
| *greatwall* |
| *pebble* |
| *exuperantia* |

**Table S1: mRNAs likely in *Drosophila* germ granules.** mRNAs with a distribution characteristic of germ granule transcripts based on *in situ* data from the Berkeley *Drosophila Genome Project* and Fly-FISH databases. Compiled based on Refs [11, 32].

| Sequence Name | Sequence (5′-3′) | Guide Score | Region targeted |
|---|---|---|---|
| *nanos* guide 1 | GTATCCAAATACATGTCCTGCAG | 0.90764 | CDS |
| *nanos* guide 2 | ATATACGACATGTTTCGCCACAC | 0.95912 | CDS |
| *nanos* guide 3 | AAGATTTTCAAGGATCGCGCAAT | 0.91676 | CDS |
| *sra* guide 1 | AACAGTAATTTGCAAGTGCACCG | 0.98239 | 5'UTR |
| *sra* guide 2 | GCATTGTCATAGTTAACACGCAG | 0.93665 | CDS |
| *sra* guide 3 | GCCAAAAGATCATGGTTGACCAG | 0.91884 | CDS |
| Direct Repeat | CAAGTAAACCCCTACCAACTGGTCGGG GTTTGAAAC | N/A | N/A |
| Terminator | TTTTTTT | N/A | N/A |

**Table S2: crRNAs.** Sequences used for crRNA arrays, generated using Cas13design [58,59].

| RNA | FW (5′-3′) | RV (5′-3′) |
|---|---|---|
| *nos crRNA* | GTATCCAAATACATGTCCTGCAG | CTTGATTGCGCGATCCTTG |
| *sra crRNA* | AACAGTAATTTGCAAGTGCACCG | CTG GTCAACCATGATCTTTTGGC |
| *nos mRNA* | GAGGAGGGCTCAACATTCTG | GCAGCAAGTGGTAGTGGTAC |
| *sra mRNA* | CCGGACCAGGACATTTTCAT | GTGGATGTTGGTCACGATGA |
| *rp49 mRNA* | CGGATCGATATGCTAAGCTGT | GCGCTTGTTCGATCCGTA |

**Table S3: qPCR primers.** Transgene specific primers used for amplification of pre-processed crRNAs, *nos* and *sra* mRNAs, and *rp49* baseline.

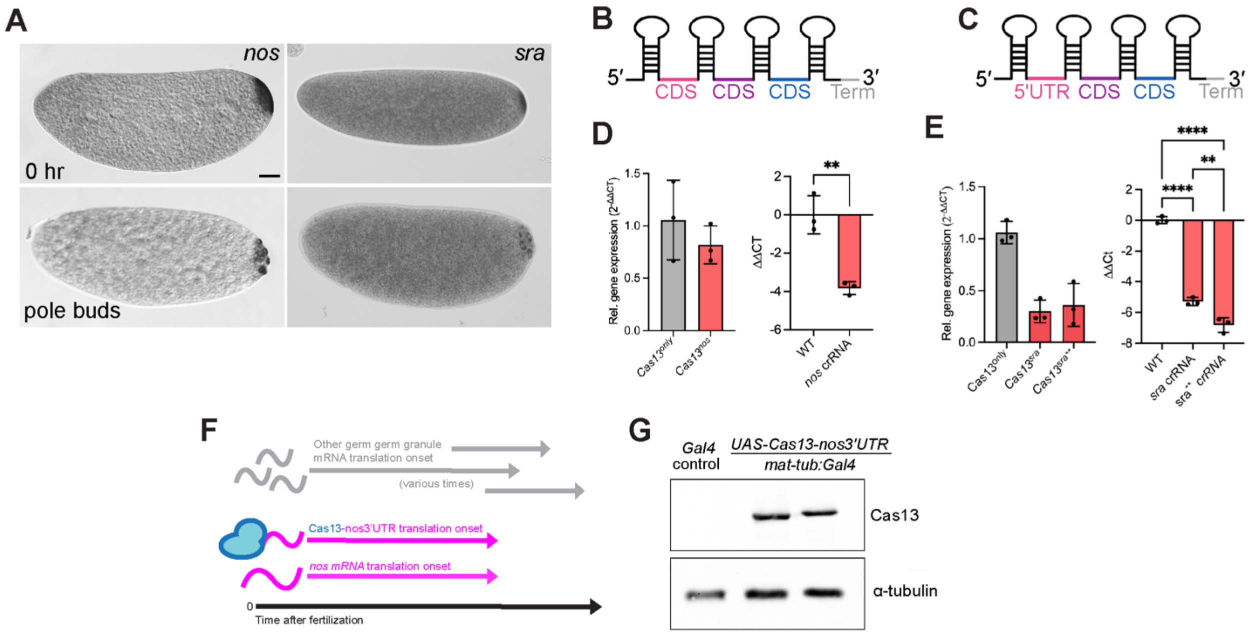


**Figure S1: System design and confirmation of transgene expression.** (A) *In situ* hybridization to detect *nanos* and *sra* mRNAs in embryos prior to and during pole cell formation. (B,C) crRNA array designs. The *nanos* crRNA array (B) includes three crRNAs targeting the coding region; the *sra* crRNA array (C) contains one crRNA targeting a region in the 5'UTR, and two crRNAs targeting the coding region. (D, E) Quantification of *nanos* (D) and *sra* (E) crRNA expression in the ovary by qPCR. Values were normalized to a wild-type control. Females with a double insertion of the *sra* crRNA array (*sra***) expressed more pre-processed crRNAs than those with a single insertion. (F) Schematic depicting translational onset of *Cas13-nanos3'UTR* transcripts. Cas13 translation will begin when *nanos* translation initiates, during late oogenesis and continues after fertilization. (G) Cas13 protein was detected in ovary extracts from two independent transgenic lines by anti-HA western blotting. **α**-tubulin was used as a loading control. Data points represent technical replicates; **$p<0.01$, ****$p<0.0001$ as determined by unpaired t-test (D) or ordinary one-way ANOVA and Tukey's multiple comparisons test (E). Scale bar = 50 µm (A).

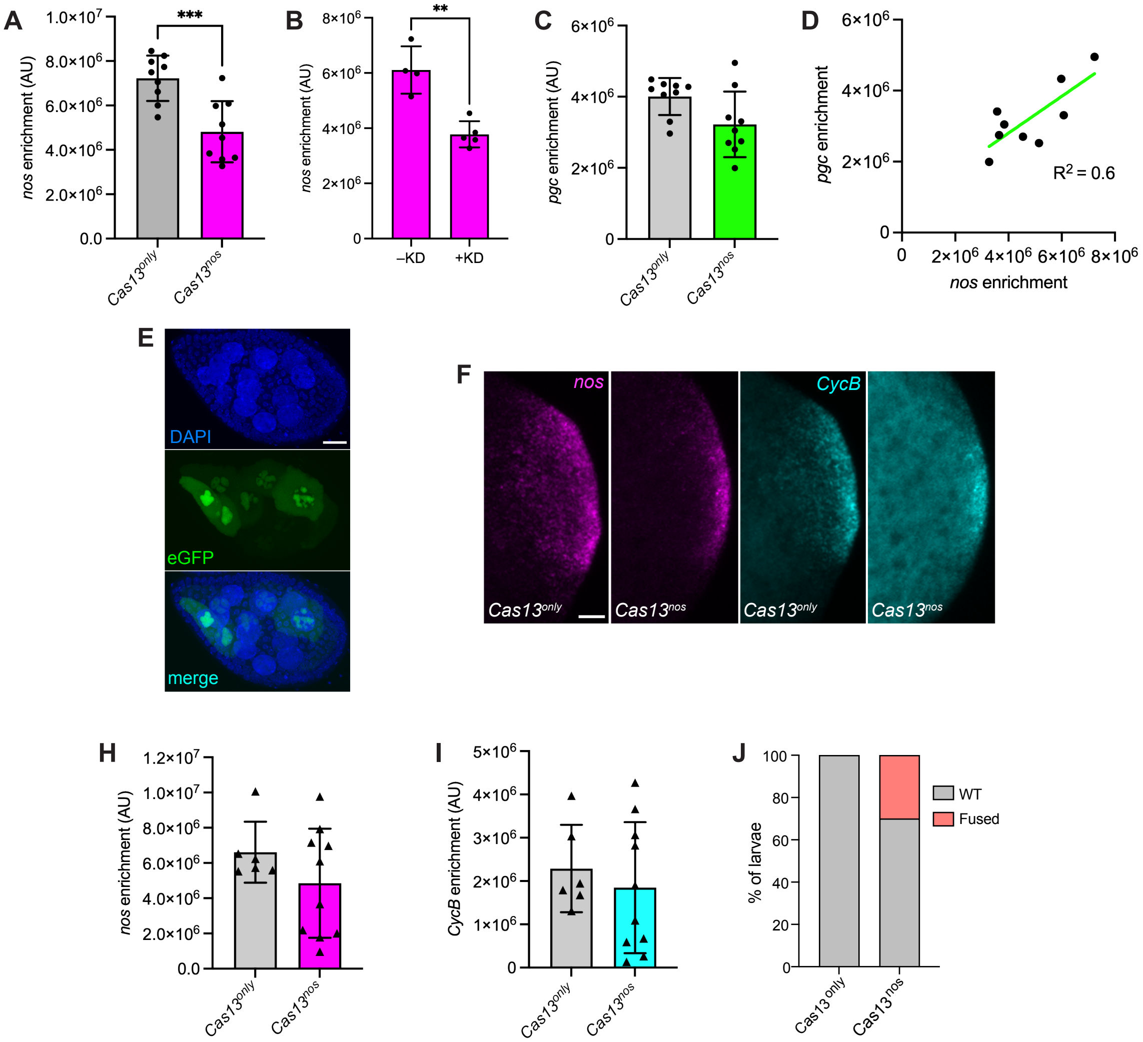


**Figure S2: Additional analysis of *Cas13^nanos^* embryos.** (A) Quantification of *nanos* enrichment for a third experimental replicate ($n$ = 8 embryos for *Cas13^only^*, $n$ = 10 embryos for *Cas13^nos^*). (B) Comparison of *nanos* enrichment between *Cas13^nos^* embryos with and without knockdown. (C) Quantification of *pgc* enrichment from the third experimental replicate. (D) Simple linear regression of the *nanos* and *pgc* enrichment for *Cas13^nos^* ($R^2 = 0.8847$, $y = 1.050x - 366335$). (E) Maximum intensity projection of stage 9 oocyte expressing a *UAS-egfp* reporter transgene driven by *mat-tub:Gal4* (green). Mosaic expression in nurse cells resulted in varying levels of transgenic mRNA produced and supplied to each oocyte. Nuclei were stained with DAPI (blue). (F) Summed slice intensity projections of embryos from a second experimental replicate of *Cas13^only^* and *Cas13^nos^* embryos with *nanos* (magenta) and *CycB* (cyan) detected by smFISH. (H, I) Quantification of enrichment of *nanos* (H) and *CycB* (I) for experiment shown in (F). (J) Quantification of larval cuticles exhibiting wild-type abdominal segmentation or fused segments ($n$ = 74 cuticles for Cas13^only^, $n$ = 76 cuticles for *Cas13^nos^*). Data points represent individual embryos. **p<0.01 as determined by an nnpaired t-test. Scale bars = 20 µm

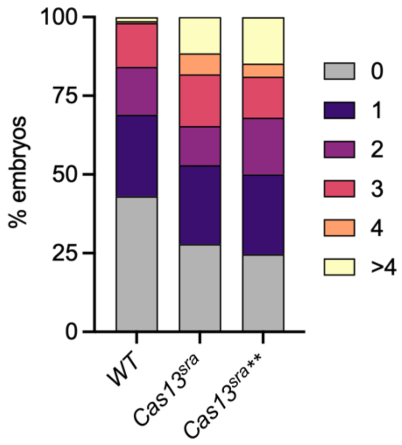


**Figure S3: Reproducibility of pole migration defects in embryos with Cas13-mediated knockdown of *sra*.** Quantification of migration defects in second experimental replicate of Cas13-mediated *sra* knockdown (*n* = 104 embryos for *Cas13$^{sra}$*, *n* = 122 embryos for *Cas13$^{sra**}$*).
